# Functional organization of the human visual system at birth and across late gestation

**DOI:** 10.1101/2025.09.22.677834

**Authors:** Vladislav Ayzenberg, Michael Arcaro

## Abstract

Understanding how the brain’s functional architecture emerges prior to substantial postnatal visual experience is crucial for determining what initial capabilities infants possess and how they learn from their environment. Using resting-state fMRI from 584 neonates in the Developing Human Connectome Project, we provide the first comprehensive systems-level characterization of human visual cortex within hours of birth and across the third trimester of gestation. We discover that newborns possess a sophisticated visual architecture already functionally organized into three distinct pathways (ventral, lateral, and dorsal), each exhibiting posterior-to-anterior hierarchical structure and adult-like topographic organization. This tripartite visual organization differs from the bipartite organization observed in macaques, suggesting this architecture emerges through intrinsic developmental mechanisms rather than being a product of extensive postnatal experience and environmental adaptation. Moreover, pathway segregation, hierarchical ordering, and connectivity maturity all strengthen progressively with gestational age, revealing that visual cortical organization emerges through an active developmental program that unfolds across late gestation. Yet, despite this large-scale structure, individual pathways follow strikingly different maturation trajectories: dorsal areas exhibit a near-adult-like functional organization, even at the earliest gestational timepoints tested, whereas ventral areas remain immature and poised for experience-dependent refinement. These findings reframe our understanding of early visual development by revealing that complex functional networks emerge before substantial visual experience, yet are differentially prepared for plasticity, providing crucial insights into how evolution has optimized the brain for rapid learning while maintaining the flexibility needed for adaptation to diverse environments.

## Introduction

Within hours of birth, human infants can discriminate basic shapes and faces (e.g., newborns recognize their mother’s face) (*1*, *2*), track motion trajectories (e.g., an object moving behind an occluder) (*3*, *4*), and categorize items by their spatial relations (e.g., above vs. below) (*5*, *6*). Newborns also rapidly learn from visual experience, forming memories and adapting their preferences based on environmental input (*7–9*). These early perceptual abilities and the capacity for rapid visual learning imply that the newborn visual system is sufficiently organized to support behaviorally meaningful processing. Understanding when this organization arises over gestation and how closely it resembles the adult state will provide critical insights into how experience shapes the brain’s architecture and what constraints this early architecture imposes on such perceptual functioning.

In adults, visual behaviors are supported by a hierarchy of cortical areas organized along ventral, lateral, and dorsal visual pathways (*10–12*). While these pathways are interconnected, each supports specialized functions: the ventral pathway is crucial for object identification (*13*, *14*), the dorsal pathway supports spatial processing and visually guided action (*10*, *11*, *15*), and a proposed lateral pathway supports dynamic social perception, such as motion-informed analysis of faces and bodies (*12*, *16*, *17*). A key organizing feature of these pathways is their topographic mapping of visual space. Adult visual areas are retinotopically organized, with adjacent neurons responding to adjacent points in the visual field, and with eccentricity (distance from the center of gaze) serving as a major organizing axis both within and across areas (*18–21*). Critically, such topographic maps can emerge from intrinsic, self-organizing developmental mechanisms even before substantial visual experience (*22–25*). These observations provide concrete predictions about the neonatal brain: that some elements of adult-like topographic organization, including pathway layout, hierarchical ordering, and large-scale functional architecture, may already be present at birth.

Yet, because most visual neuroscience has focused on older children or animal models, we still lack a clear account of how the human visual system is organized at birth. A few months, or even a few hours, of postnatal experience can substantially alter cortical organization (*1*, *26–28*), underscoring the importance of examining human brain structure prior to extensive visual input. Moreover, most non-human mammals are more precocial than human infants, with faster neural development and earlier maturity, (*29–31*), raising questions about the extent to which findings from other newborn animals generalize to human infants (*32*). Human visual cortex also undergoes substantial changes extending into adolescence (*33–36*), a trajectory that may support greater flexibility and adaptation than is observed in other species (*37*, *38*). Finally, developmental studies typically examine individual or small groups of regions in isolation (*39–42*), providing only limited insight into how pathways develop relative to one another.

The lack of data from human neonates limits our understanding of how ontogenetic and phylogenetic factors contribute to the structure of the human brain. Adult humans and non-human primates exhibit substantial differences in visual system organization. Whereas macaques show evidence of two visual pathways, ventral and dorsal, humans show evidence of a third lateral pathway specialized for social perception. Two competing hypotheses could explain this species difference. First, the lateral pathway may arise during human development through extensive experience with complex social dynamics (*43*). The prolonged developmental trajectory of the human visual system may support such plasticity (*33–36*), and may explain why the cortical expansion of lateral areas between infancy and adulthood mirrors the expansion observed between macaque monkeys and human adults (*44*). Under this developmental hypothesis, human ontogeny recapitulates phylogeny such that the infant visual system initially contains a two-pathway structure like macaques, but, develops a distinct lateral pathway through postnatal experience. Alternatively, the lateral pathway may represent a human-specific phylogenetic innovation already present at birth. The medial shift of V1 in humans compared to its lateral position in monkeys has allowed for lateral expansion of the visual system (*45*), which may accommodate a dedicated social pathway. However, because visual cortical organization has been primarily characterized in adults or older children, the relative contributions of environmental and evolutionary factors to human visual cortical development remains unknown.

In the current study, we directly tested these competing developmental and evolutionary hypotheses by characterizing the large-scale organization of human visual cortex at birth. To accomplish this, we leveraged a large resting-state fMRI dataset from the Developing Human Connectome Project (DHCP) (*46*) spanning a broad range of gestational ages (26 to 46 weeks), including infants scanned within hours of birth. This dataset enabled us to chart how visual cortical organization unfolds during late gestation and to directly test whether human-specific pathway architecture emerges through intrinsic developmental mechanisms or requires postnatal experience. Using a surface-based atlas of adult retinotopic visual areas, we assessed signatures of large-scale organization in neonates, including processing pathway segregation, hierarchical structure, areal boundaries, and spatial topography. We quantified the developmental maturity of each cortical area by comparing neonatal connectivity profiles to those of young adults. Together, these analyses provide the first comprehensive, systems-level account of human visual cortical architecture at birth and reveal how distinct aspects of organization emerge across gestational development.

## Results

### Organization of the human visual cortex at birth

To investigate the functional organization of the visual cortex at birth, we analyzed time-varying BOLD signals from neonates scanned on their first day of life. Functional correlations were computed between 24 cortical visual areas (48 across both hemispheres) defined by projecting an adult probabilistic atlas onto each neonate’s cortical surface and high-resolution anatomical volume image (see Methods; Supplemental Figure 1). We then assessed whether neonatal visual cortex exhibited adult-like organization in terms of area boundaries, visual pathway grouping, hierarchical ordering, and functional fingerprint patterns.

### Arealization of the neonate visual cortex

We first examined whether neonate visual cortex shows evidence of arealization, defined as the presence of distinct functional correlation patterns that reliably differentiate each area from its neighbors. To do so, we computed seed-to-whole-brain correlations between the mean timeseries in each area with the time series of each voxel in the contralateral hemisphere. Qualitative examination of these correlation maps revealed that correlations for each area generally peaked in their cross-hemisphere homolog (e.g., left V1v to right V1v) at the group level (Figure 1A). Importantly, homotopic correlations were also quantitively stronger between homotopic areas (e.g., left V1v and right V1v; F(2, 78) = 493.06, *p* < .001, 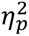 = 0.93) than between adjacent (e.g., left V1v and right V1d; *p* < .001, *d* = 0.41, BF_10_ > 100) and with distal regions (e.g., left V1v and right MT; *p* < .001, *d* = 1.9, BF_10_ > 100), in neonates (Figure 1B). This pattern was observed across nearly all areas, with stronger homotopic correlations in every comparison relative to distal areas, and all but three (hV4, PHC1, LO2) also stronger than adjacent areas (Figure 2C). These findings suggest that areas are already functionally distinct at birth.

**Figure 1.**
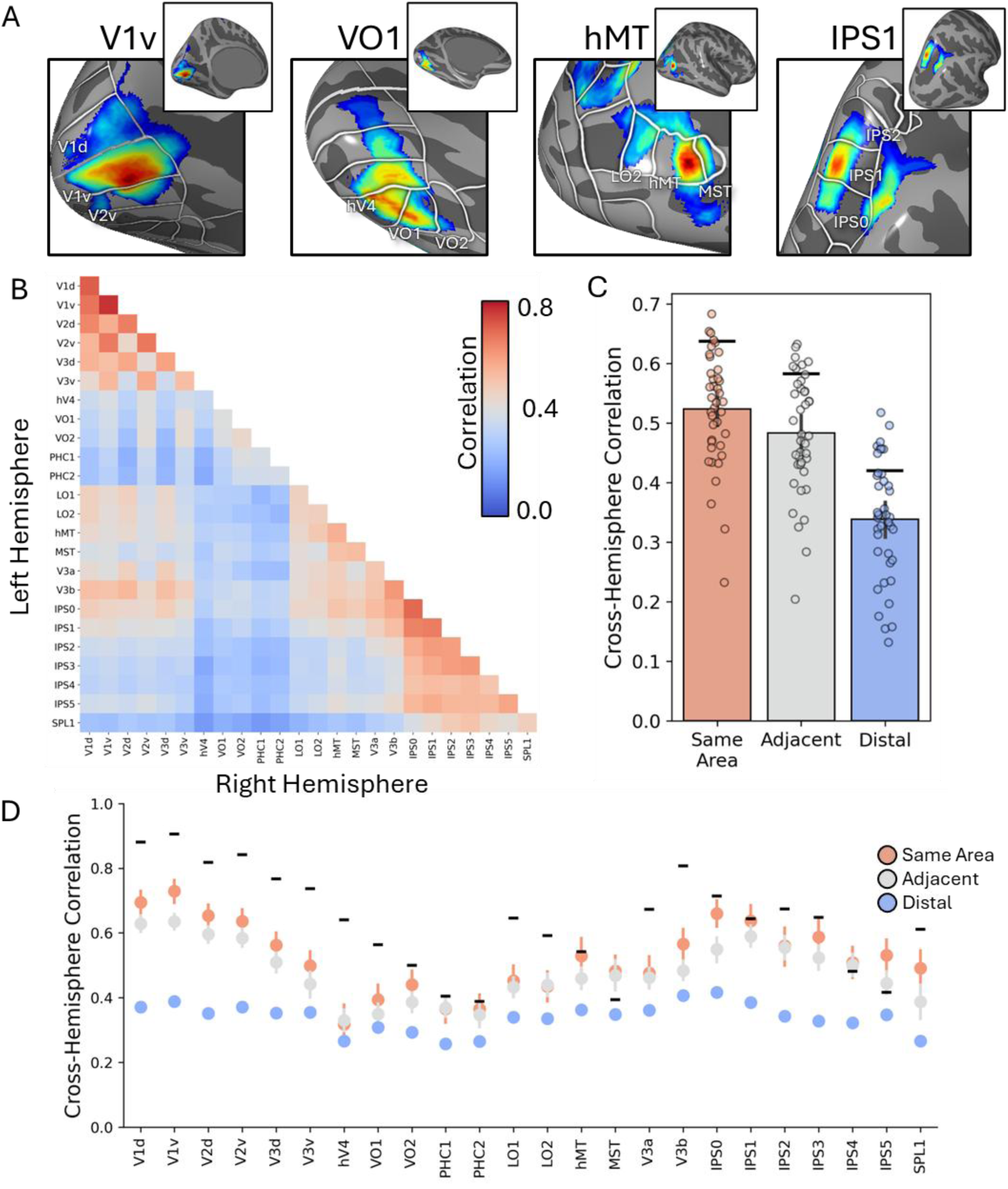
Arealization of neonate visual cortex. (A) Across-hemisphere areal correlation maps for V1v, VO1, MT, and IPS1. (B) Matrix of time series correlations between visual areas in the left and right hemispheres. (C) Mean cross-hemisphere correlation (Pearson’s r) between the homotopic, adjacent, or distal visual areas. Each dot represents a neonate. (D) Cross-hemisphere correlations for each visual area with homotopic, adjacent, or distal areas in the contralateral hemisphere. Black bars represent the homotopic correlation in adults, which serve as a noise ceiling. Dots correspond to individual participants.

**Figure 2.**
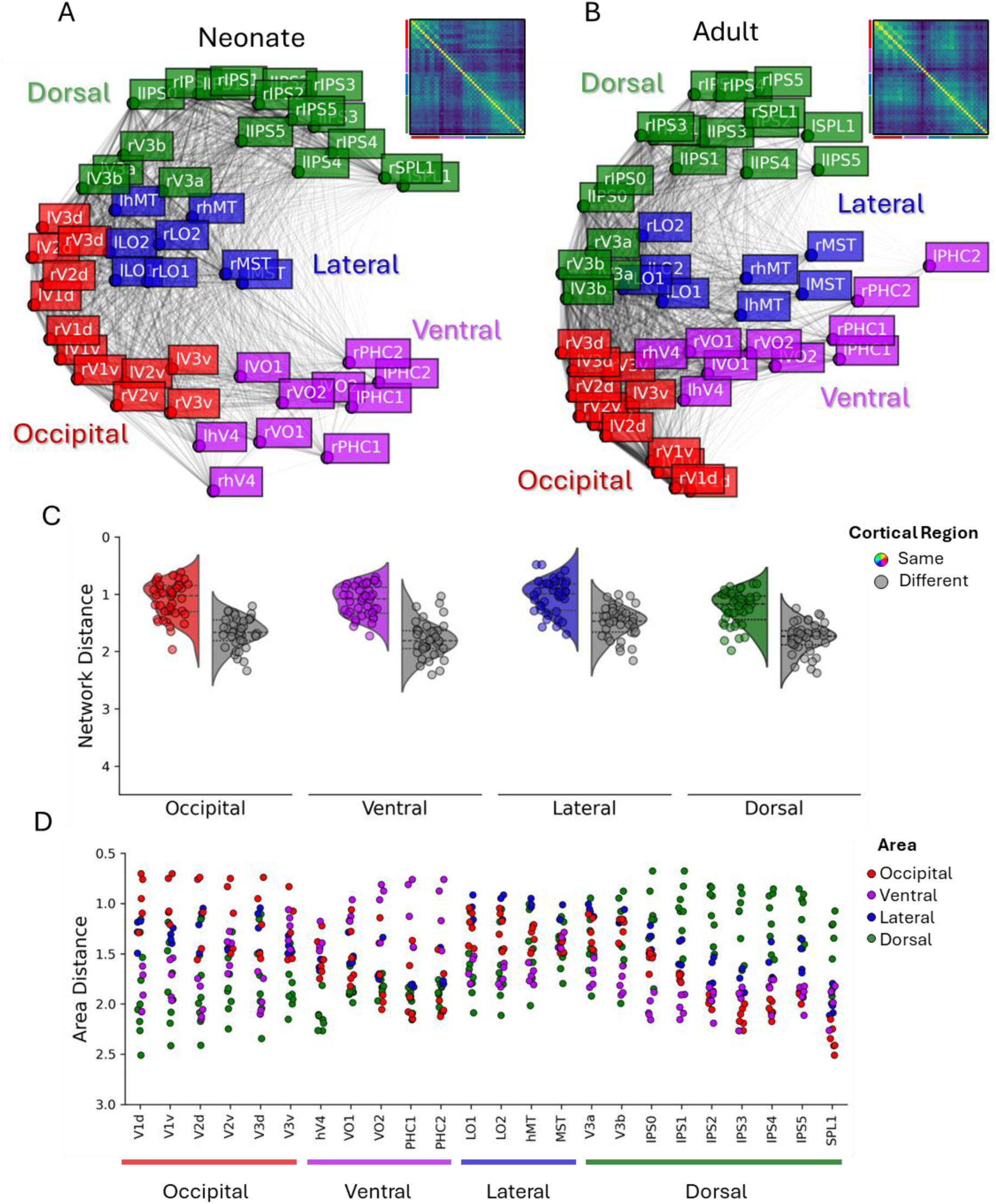
Hierarchical organization of visual cortical areas in neonates. (A-B) MDS projection of time series correlations among visual cortical areas in (A) neonates scanned on the first day of life and (B) adults. Insets show full pairwise correlation matrices. The colored lines correspond to (red) occipital, (purple) ventral, (blue) lateral, and (green) dorsal areas. (C) Distance in MDS space between areas in the same (colored) or different (gray) pathway groupings. Data for both same and different networks was averaged across areas. Dots correspond to individual participants. (D) Distances from each area to all other areas, color coded by pathway grouping (C-D) The Y-axis is inverted for ease of viewing, with smaller distances towards the top.

### Pathway organization and hierarchical structure in the neonate visual system

A defining feature of visual cortical organization is its separation into three anatomically and functionally distinct pathways, ventral, lateral, and dorsal (*12*, *17*, *47*, *48*), each of which originates in posterior occipital cortex and progresses anteriorly, forming a hierarchical structure that supports increasingly complex visual processing. To what extent does the functional organization of neonate visual areas correspond to the anatomical segregation of areas into occipital, ventral, lateral, and dorsal networks?

To assess whether this large-scale organization is present at birth, we used multi-dimensional scaling (MDS) to project interareal correlations of the pairwise time series between all visual areas into a two-dimensional space (Figure 2A). The MDS projections revealed clear separation of visual areas into four distinct groupings: occipital (V1 through V3), ventral, lateral, and dorsal cortices. This pattern closely resembled that of adults (Figure 2B), with the spatial layout of neonatal areas significantly correlated with adults (*r* = 0.96, *p* < .001). Further, the distances between areas in the MDS projection were significantly smaller between areas in the same anatomical grouping than across groupings, (F(1, 39) = 319.47, *p* < .001, 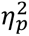 = 0.89). Post-hoc comparisons confirmed this pattern for all four groupings (*ps* < .001; *ds* > 1.01; BF_10_ > 100; Figure 2C) and at the level of individual area level where most areas had their nearest neighbor within the same group (Figure 2D). These results held when the MDS projection was restricted to cross-hemisphere correlations, ruling out effects of spatial autocorrelation and local signal smoothing from preprocessing steps (see Supplemental Figure 2).

Next, we examined whether, the spatial layout of cortical visual areas also reflected the posterior-to-anterior hierarchical ordering typically observed in adults. The MDS projection showed orderly progressions from early visual areas in the occipital cortex (V1 to V3), to intermediate and anterior regions in each pathway (hV4 through PHC2 in the ventral pathway, from LO1 to MST in the lateral pathway, and from V3A/B through SPL1 in the dorsal pathway; Figure 2A). To quantify this, we compared the MDS-derived order of areas to a prototypical posterior-to-anterior hierarchy defined based on anatomical location and prior adult studies (*48*). Across neonates, there was a significant correlation between the prototypical hierarchical order and the MDS-derived arrangement for all pathway groupings: occipital (*r*_mean_= 0.49; 95% CI [0.26, 0.71]), ventral (*r*_mean_= 0.26; 95% CI [0.13, 0.38]), lateral (*r*_mean_= 0.23; 95% CI [0.04, 0.40]), and dorsal cortices (*r*_mean_= 0.41; 95% CI [0.28, 0.51]). These patterns were also observed in the adult data for occipital (*r*_mean_= 0.41; 95% CI [0.32, 0.52]), ventral (*r*_mean_= 0.59; 95% CI [0.52, 0.65]), lateral (*r*_mean_= 0.25; 95% CI [0.16, 0.34]), and dorsal (*r*_mean_= 0.53; 95% CI [0.48, 0.57] pathway groupings. These results indicate that the neonatal visual system already exhibits adult-like grouping and an emerging hierarchical structure at birth

### Maturity of visual cortical areas

Having established that the large-scale organization of the visual cortex is already in place at birth, we next assessed the functional maturity of visual pathways and individual areas. In the mature visual system, each visual pathway supports distinct functions, and each area contributes uniquely to information processing. If these areas already have functional differentiation at birth, we would expect visual pathways and their constituent areas to exhibit adult-like patterns of functional activity. To test this, we compared the *functional fingerprint* of each visual area in neonates to its counterpart in adults. Each functional fingerprint was defined as its pattern of temporal correlations with all other areas (see Methods), derived from the previously computed ROI-to-ROI temporal correlation matrices (see Figure 2A). This approach builds on the idea that cortical areas possess distinct structural connectivity profiles (*49*), which will give rise to distinct patterns of functional correlations. By comparing neonatal and adult fingerprints, we assess the extent to which each area expresses a mature pattern of functional organization at birth.

We first evaluated the specificity of each neonate area’s functional fingerprint by comparing it to the fingerprints from all areas in the adult brain. If neonatal areas already exhibit adult-like specialization, we expect each area’s fingerprint pattern to be most similar to its adult counterpart. We tested this at two levels of granularity: cortical region and individual areas. At the regional level, we asked whether areas within a given cortical region in neonates were more similar to adult areas from the same region than to those from different cortical region. This analysis revealed a main effect of cortical region grouping, with neonate areas showing significantly stronger correlations with adult areas within the same region grouping (e.g., neonate V1 to adult V1) than from other region groupings (e.g., neonate V1 to adult MT; Figure 4A) (*F*(1,29) = 808.43, *p* < .001, 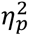 = .51). Post-hoc comparisons (Holms-Bonferroni corrected) confirmed this pattern for all neonate regional groupings (Figure 3A; Occipital: *d* = 3.48; Ventral: *d* = 1.07; Lateral: *d* = 2.65; Dorsal: *d* = 5.24; *ps* < .001). At the level of individual areas, we asked whether each neonate area was most similar to its adult counterpart, regardless of cortical region. In most cases, the functional fingerprint of a neonate area was most strongly correlated with the same area in adults, and typically ranked among the top three correlations across all possible pairings (e.g., neonate V1 with adult V2; Figure 3C).

**Figure 3.**
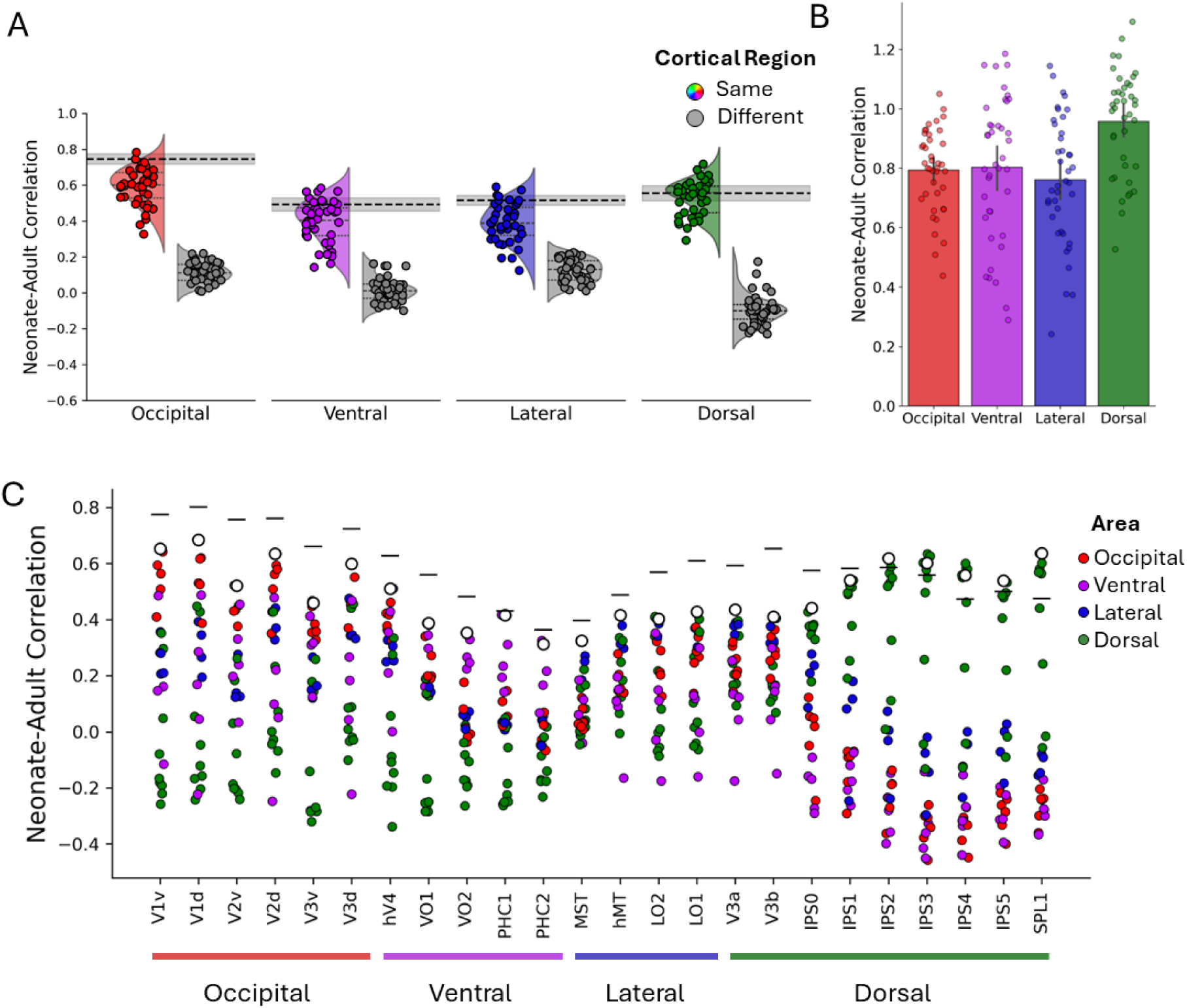
Maturity of cortical visual connections. (A) Pathway-level similarity between neonates and adults for occipital, ventral, lateral, and dorsal visual areas. Colored distributions show the correlation between visual areas in neonates with the same network in adults. Gray distributions show the correlation with different networks. The dotted black line with shading depicts the adult noise ceiling. Data for both same and different cortical regions was averaged across areas. (B) Fingerprint similarity normalized by the adult noise ceiling for occipital, ventral, lateral, and dorsal pathways. (A-B) Dots correspond to individual participants. (C) Area-level similarity between neonatal and adult connectivity fingerprints. White dots indicate the correlation with the corresponding area in adults and colored dots indicate the correlations with other visual areas. Visual areas are color coded by anatomical cluster. Black lines depict the adult noise ceiling for each region.

**Figure 4.**
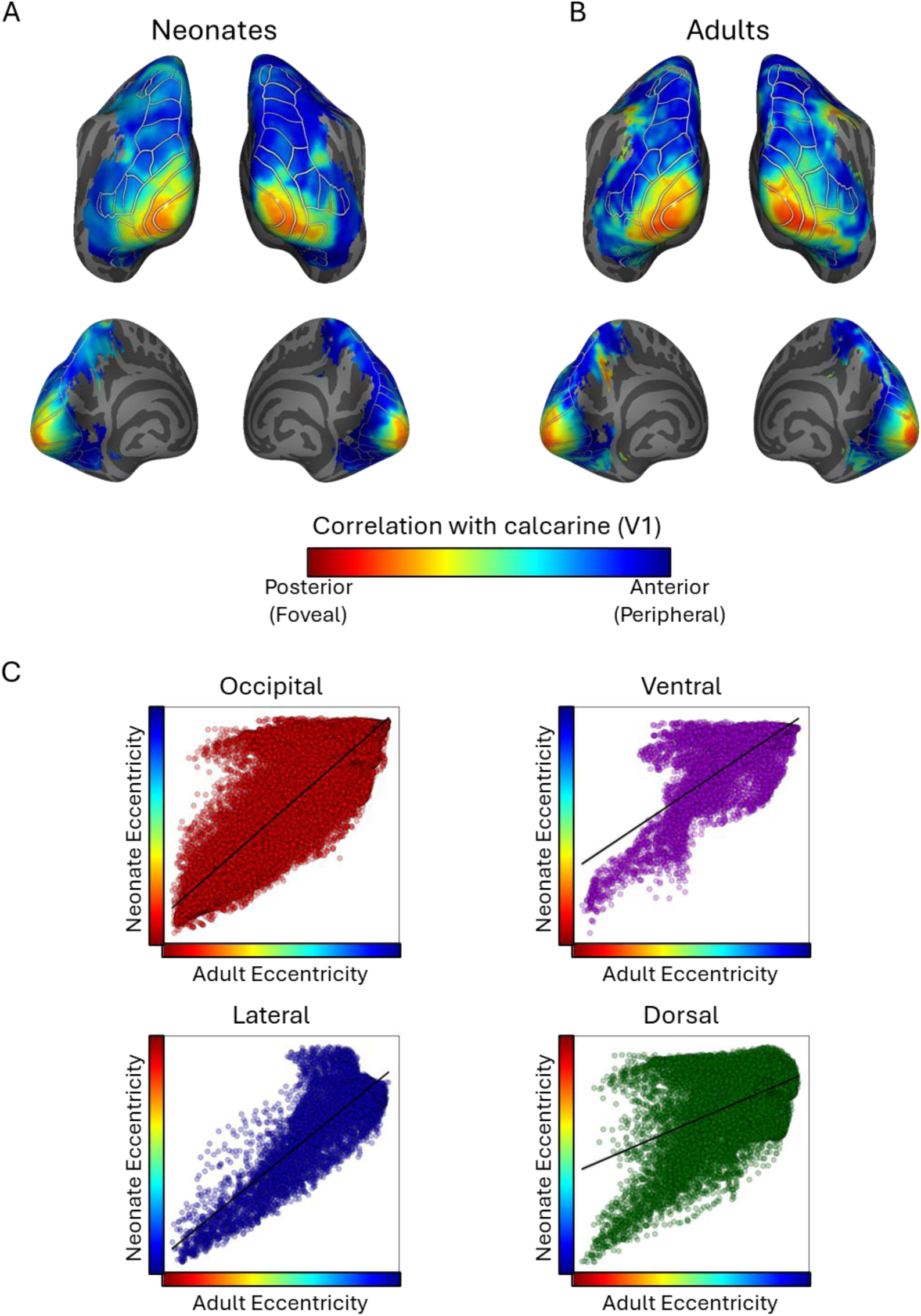
Eccentricity organization in the neonate visual system. (A-B) Foveal-to-peripheral sensitivity maps showing the relative voxel-wise sensitivity to ROI bands arranged posterior-to-anterior along the calcarine sulcus in (A) neonates and (B) adults. Red values correspond to the most foveal ROI band, whereas bluer values correspond to more peripheral ROI bands. Eccentricity maps are plotted within an extended visual system ROI. (C) Scatter plots illustrating the correlation between eccentricity values for neonates and adults.

We then assessed the relative maturity of each cortical region by comparing neonate-adult fingerprint correlations to the adult noise ceiling, which estimates the upper bound of similarity expected across adults. The dorsal pathway was the only group with a mean neonate-adult correlation that fell within the adult noise-ceiling confidence intervals. A majority of neonates exceeded the lower bound of the adult noise ceiling for the dorsal areas (26/40), but not for occipital (3/40), ventral (14/40), or lateral areas (10/40). When neonate-to-adult correlations were normalized by the adult noise ceiling, we found a significant main effect of regional grouping (*F*(3, 87) = 58.85, *p* < .001, 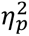 = .67). Post-hoc comparisons (Holms-Bonferroni corrected) showed that dorsal areas had significantly higher normalized fingerprint correlations than all other pathway groupings (*ps* < .001, *ds* > .73; BF_10_ > 100). While the large-scale organization of the neonate visual system is largely adult-like, these findings indicate that the maturity of visual areas varies across pathways, with the dorsal pathway showing the most adult-like pattern of interareal correlations at birth.

### Topographic organization of visual space in neonates

A fundamental organizing principle of the visual system is the existence of a continuous representation of stimulus eccentricity in all visual pathways (*19–21*, *50*, *51*). To investigate whether neonates already show evidence of an eccentricity organization, we examined the strength of cross-hemisphere voxel-wise correlations in each pathway to 5 ROI bands spanning the length of the V1 calcarine sulcus (see Methods and Supplemental Figure 3). The most posterior band corresponds to foveal cortex, and therefore small eccentricities, and more anterior bands correspond to increasingly more peripheral portions of space, and therefore larger eccentricities.

Qualitative inspection of the neonatal maps showed an adult-like eccentricity patterns across the visual system which recapitulated classic foveal vs. peripheral sensitivities of each pathway (*18*, *51–53*). Occipital areas showed greater foveal representations in posterior-lateral visual areas, and increasingly more peripheral representations in anterior-medial areas (*54*). Ventral areas exhibited the classic a lateral-to-medial organization (*18*, *51*), with lateral portions showing stronger foveal sensitivity. Like prior work (*55*, *56*), the lateral pathway showed an inferior-to-superior organization, with inferior portions being more foveally biased and superior portions being more peripherally biased. Finally, the dorsal pathway exhibited greater peripheral representations compared to the other pathway, consistent with prior work showing stronger peripheral input to the dorsal pathway and larger receptive fields (*52*).

The qualitative similarity between neonates and adults was also found quantitatively, with significant correlations between the voxel-wise eccentricity responses of every pathway between neonates and adults (occipital: *r =* 0.85, 95% CI [0.85, 0.86]; ventral: *r* = 0.74, 95% CI [0.73, 0.75]; lateral: *r* = 0.84, 95% CI [0.83, 0.84]; dorsal: *r* = 0.49, 95% CI [0.47, 0.50]). Thus, the neonate visual system may already exhibit an adult-like eccentricity organization.

### Development of the visual system across gestational time

The preceding analyses demonstrated that, by full-term birth, the visual system is functionally organized into distinct pathways that vary in their degree of maturity. To better understand how this architecture unfolds during development, we next examined visual cortical organization in neonates born at four gestational ages: pre-term (26–32 weeks), early-term (32–38 weeks), term (38–42 weeks), and post-term (42–46 weeks).

We first asked whether the large-scale spatial layout of visual pathways becomes more adult-like with increasing gestational age. MDS projections of interareal dissimilarity revealed a progressive developmental trajectory. Neonates born at term and post-term showed a clear grouping of visual areas into occipital, ventral, lateral, and dorsal cortices (Figure 5A), resembling the adult pattern (Figure 2B). Within each regional grouping, areas appeared hierarchically arranged from early visual areas in occipital cortex to intermediate and then anterior regions along each processing pathway. Early-term neonates also showed pathway-level organization, particularly along the dorsal pathway, but grouping of occipital, posterior lateral, and ventral visual areas were less distinct (Figure 5A). In contrast, pre-term neonates showed a more diffuse spatial layout, with overall weaker separation of visual areas into coherent pathway groupings. Notably, however, the dorsal pathway appeared hierarchically structured even in this youngest group, suggesting that this organizational scaffold may emerge earlier than other pathways.

**Figure 5.**
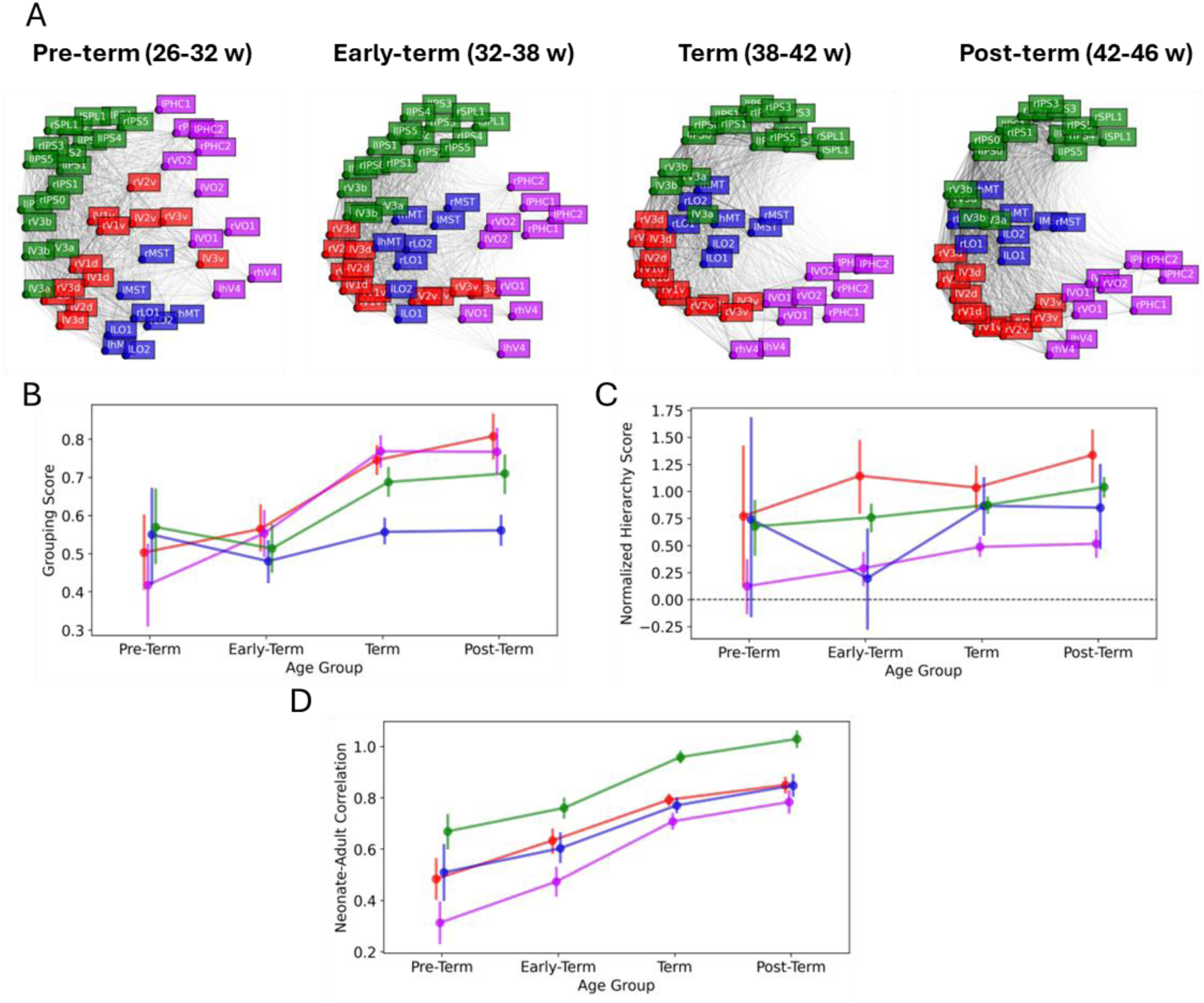
Development of the visual system across gestational age. (A) MDS projections of interareal dissimilarity matrices, derived from time series correlations between visual cortical areas, for neonates born pre-term, early-term, term, or post-term. (B) Grouping score as a function of gestational age indicating the degree to which visual areas group according to their anatomically defined network. The grouping score is computed as the distance between areas in different networks subtracted from the distance between areas in the same network. (C) Normalized hierarchy score as a function of gestational age indicating the degree to which visual areas follow the anatomically defined ordered arrangement of areas in each visual network. The normalized hierarchy score is computed as the correlation between the anatomical order of areas for a network with the order computed using MDS projections for each neonate. (D) Maturity as a function of gestational age as indicated by the correlation between fingerprint connectivity for a region in individual neonates with the same regions in adults. To evaluate the relative maturity of each correlation, neonate correlations were normalized by dividing them by the adult correlation for a visual region.

We quantified these developmental changes across three aspects of visual cortical organization: visual pathway grouping, hierarchical ordering, and regional-level maturity. First, we measured the degree to which areas clustered into known visual pathways. Grouping strength increased with gestational age, with term and post-term neonates showing the clearest segregation of visual pathways (*F*(3, 580) = 24.62, *p* < .001, 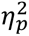 = .11) (Figure 5B). Second, we assessed whether the ordering of areas within each pathway followed the expected hierarchical progression from early visual areas in occipital cortex to posterior and then anterior areas of each pathway. Pre-term neonates lacked reliable hierarchical ordering along ventral and lateral pathways (*ps* > .129; *ds* < 0.29), and early-term neonates showed no hierarchical ordering in the lateral pathway (*p* = 0.385, *d* = 0.09). In contrast, hierarchical ordering was present in occipital (*ps* < .025; *ds* > 0.44) and dorsal (*ps* < .001; *ds* > 0.97) pathways for all age groups with the dorsal pathway showing the strongest effect size at each age (Figure 5C). Third, we assessed the degree of functional maturity of each cortical region by measuring the similarity of connectivity fingerprints between neonates and adults. Fingerprint maturity increased linearly with gestational age across all regions (*F*(3, 580) = 76.74, *p* < .001, 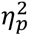 = .28), with dorsal areas consistently showing the highest maturity at all age groups (*ps* < .025, *ds* > 0.64), and ventral areas the lowest at all ages groups (*ps* < .020, *ds* > 0.26) (Figure 5D). Together these results suggest that the large-scale organization of the visual system emerges and becomes progressively more adult-like over gestation.

## General Discussion

In this study we characterized the functional organization of the human neonatal visual system. We found that by full-term birth (at least 38-weeks gestation), the visual cortex is already organized into three functionally differentiated pathways (ventral, lateral, and dorsal) that resemble adult visual networks. Each pathway originates in occipital cortex and progresses anteriorly, forming a posterior-to-anterior hierarchy of areas. Importantly, although the overall scaffold is present at birth, pathways differed in their degree of maturity, with dorsal regions showing more adult-like connectivity than the others. This large-scale architecture becomes progressively more defined with gestational age up to full-term birth. These results demonstrate that human infants begin life with a visual system that is broadly adult-like in its large-scale architecture, yet marked by pathway-specific developmental trajectories that continue to unfold after birth.

### Gestational emergence of large-scale visual pathway organization

Our finding that visual pathway organization becomes progressively more defined across gestational development builds on established principles of staged cortical differentiation *in utero*. Prior fetal imaging and postmortem studies demonstrate that broad regional distinctions in cortical morphology and progenitor populations are present by the end of the first trimester (*57–59*), whereas differentiated areal identity, including region-specific transcriptional signatures, cytoarchitecture, and afferent connectivity, emerges later, around 22 weeks gestation (*58–60*). By this stage, visually evoked responses can be detected in preterm infants (*61*), yet microstructural differentiation (*58*, *62*), cortical folding (*59*, *63*, *64*), and interareal connectivity (*60*, *65*, *66*) continue to develop throughout the third trimester. Our results extend this literature by showing that functional organization is relatively diffuse in preterm and early-term human neonates, with weaker segregation of areas and less reliable hierarchical ordering, but becomes more coherent by term age. This suggests that fetal and preterm differentiation provides a provisional scaffold that is refined into an adult-like pathway-level architecture during late gestation.

Thalamocortical connectivity likely contributes to this transition from scaffold to functional networks (*67*). Prior anatomical and diffusion imaging studies show that thalamic afferents, particularly from the pulvinar and LGN, mature during later gestation (*68–70*). Our own diffusion work show that pulvino-cortical structural connections to dorsal cortex mature earlier than those to ventral and lateral regions (*71*), a pattern consistent with the advanced maturity of the dorsal pathway observed in the present study. In contrast, occipital cortex shows relatively early structural connectivity with the thalamus but more protracted functional organization, suggesting that intracortical refinement and visual experience are critical for its full maturation (*72*, *73*). Taken together, these findings are consistent with the possibility that thalamocortical inputs bias the timing of cortical development, providing an early scaffold for dorsal processing, while occipital and ventral systems may rely more heavily on later-developing intracortical mechanisms and postnatal input.

### Pathway-specific maturation in human visual development

Our functional fingerprint analysis revealed distinct developmental trajectories across cortical regions and pathways. This aligns with prior developmental work, which has largely focused on individual areas or small clusters of cortical regions in older children and non-human animals (*32*, *39–42*, *74*). For example, V1 shows the earliest signatures of maturity, but continues to refine its microarchitecture, connectivity, and functional tuning after birth (*72*, *73*, *75*, *76*). Orientation selectivity sharpens with age (*72*, *77*, *78*) and can be disrupted by deprivation (Blakemore & Cooper, 1970), while ocular dominance can be altered by monocular deprivation (*79–81*). Similarly, area MT exhibits early cytoarchitectural and functional maturation (*82*), but undergoes reorganization of its connectivity. In marmosets, MT initially receives strong input from the pulvinar at birth, but over time this subcortical input is reduced (though not lost), and V1 becomes its primary source of input (*76*, *83*). Human infant fMRI (5- to 8-week olds) likewise reveals immature V1–MT coupling (*84*, *85*). These examples illustrate staggered area-specific trajectories. Finally, existing research shows that the ventral areas follow a more protracted trajectory compared to other pathways, remaining highly sensitive to postnatal input, with specialization shaped by experience with faces, objects, and culturally specific stimuli such as written words (*86–88*).

Our findings indicate that the dorsal pathway shows more adult-like connectivity patterns at birth than other visual pathways. This early maturation is consistent with evidence from nonhuman primates and humans showing early cytoarchitectural differentiation of posterior parietal cortex (*76*, *89*) and earlier development of magnocellular compared to parvocellular inputs (*90*, *91*). Such early maturation has functional consequences: dorsal areas exhibit reduced capacity for reorganization after injury and less plasticity across childhood compared to ventral cortex (*92*, *93*). Prenatal disruptions to dorsal areas are likewise associated with selective cognitive and perceptual impairments (*94*), consistent with the dorsal vulnerability hypothesis (*95*). Taken tougher, these converging lines of evidence suggest that the relatively early maturation of the dorsal pathway constrains its later plasticity. Our results extend this body of work by showing that the dorsal pathway, at a systems level, reaches relative maturity earlier, providing infants with scaffolds for spatial orienting, motion processing, and attentional control. Whereas prior work has primarily emphasized a posterior-to-anterior sequence of maturation (*96–98*). Our findings extend this principle by showing that development also differs across pathways, with dorsal cortex achieving earlier functional maturity than ventral cortex (cf. Bourne et al. (*99*).

### Human-specific lateral pathway differentiation at birth

Our results also reveal a robust tripartite organization in full-term human neonates, distinguishing ventral, lateral, and dorsal pathways. This tripartite motif mirrors the architecture observed in adults (*12*, *47*), and suggests that lateral pathway differentiation emerges through intrinsic developmental mechanisms rather than being solely experience-driven. In contrast, neonatal macaques exhibit only a bipartite division into dorsal and ventral systems (*32*), mirroring the adult organization (*100*, *101*). This species difference does not reflect an increase in the number of human visual areas (*102*), but rather a reconfiguration of areal relationships, with MT and neighboring regions more strongly differentiated from the dorsal parietal cortex in humans. Such reorganization may have arisen as a developmental consequence of broader evolutionary changes, including the medial shift of V1 in humans (*45*) and the expansion of lateral temporal and parietal cortices (*103*), that may have altered connectivity patterns between MT, dorsal parietal cortex, and adjacent lateral cortex. Cross-species comparisons therefore suggest that the tripartite organization visible at birth in humans reflects how evolutionary and developmental pressures have shaped the unique functional architecture of the human visual system.

### Behavioral implications of cortical development

The earlier maturation of the dorsal pathway raises the question of what functions they support in infancy. One possibility is that the dorsal pathway provides the foundation for an infant’s initial understanding of the physical world. In adults, the dorsal pathway supports a wide range of perceptual and cognitive capacities, including visual attention (*104*), spatial and relational reasoning (*11*, *105*, *106*), numerical cognition (*107*, *108*), and intuitive physical inference (*109*). Notably, several of these abilities emerge within the first months in infancy (*6*, *110–113*), suggesting that early-developing dorsal circuits may provide infants with a framework for modeling for how objects and agents interact with the physical world (*110*, *113*).

A complementary possibility is that dorsal pathway functions also serve as prerequisites for the development of other cortical networks and associated behaviors. For example, the ability to attend to and individuate objects, typically attributed to the dorsal pathway, may need to precede identifying objects based on featural information, a function more closely linked to the ventral pathway (*114–118*). This developmental ordering of behaviors is supported by evidence that spatial reasoning and object individuation emerge earlier than the ability to use featural information to identify objects (*115*, *119–121*). Rather than developing independently, dorsal and ventral functions may unfold along interdependent trajectories, with early-developing dorsal computations facilitating the emergence of ventral visual functions.

## Conclusion

Our findings provide a systems-level account of human visual cortex at birth. By characterizing the large-scale layout of visual pathways and their relative maturity across gestational age, this work bridges prenatal anatomical development with the onset of visual experience. The discovery that distinct cortical networks are already in place, yet follow different developmental trajectories, opens new avenues for investigating how postnatal experience interacts with early cortical structure to shape functional specialization. Critically, our analyses were based on spontaneous activity in sleeping neonates, which offers insight into intrinsic architecture but cannot address the functional roles of these areas during perception and behavior. Functional development can be assessed in many ways, including task-based imaging in awake infants (*122*), behavioral paradigms, and longitudinal tracking of brain–behavior relationships. Future studies combining these approaches will be essential for understanding how early connectivity patterns give rise to the dynamic and flexible visual abilities that emerge over infancy.

## Methods

### Participants

We analyzed all eligible neonate scan sessions from the Developing Human Connectome Project (dHCP) dataset (*N* = 736; 46% female; gestational age at scan = 26 to 48 weeks; age at scan = 0 to 26 days). Eligibility was defined as all neonates scanned postnatally who contributed both functional (*46*) and structural (*70*) MRI data. We excluded neonates using the following criteria, in order: (1) atlas registration failure (*n* = 35), (2) poor atlas alignment upon visual inspection (*n* =111; see *Identification of Cortical Visual Areas*), and finally (3) poor signal quality in the functional scans (after *n* = 6; see *Signal Quality*) leaving a total of 584 usable neonate scans. Neonate data were further split into five groups based on whether they contributed MRI data within the first day of life (*n* = 40; term or post-term infants only), or based on their gestational age at time of scan (pre-term 26-32 weeks: *n* = 29; early-term [32-38 weeks]: *n* = 98; term [38-42 weeks]: *n* = 304; post-term [42-46 weeks]: *n* = 153). Gestational age bins were determined using the standard classification as defined by the Centers for Disease Control (CDC) (*123*). For comparison, we also analyzed all eligible adults from the 7T Human Connectome Project (*N =* 165; 60% female; ages = 22 to 25 years). One subject was excluded for failed registration. Eligibility criteria were the same as neonates. After applying exclusion criteria, signal quality was matched across neonates of different gestational ages and adults.

### Data acquisition & initial preprocessing

Neonate functional and structural MRI data were collected in a 3T Philips Achieva scanner with a dedicated 32-channel neonatal head coil and 80 mT/m gradients (*70*, *124*, *125*). Resting-state functional images were collected using a high temporal (TR = 392 ms) multiband (factor=9) single-shot echo planar (EPI) sequence (*46*) at a resolution of 2.15 mm isotropic for 15 minutes (*124*). T1w and T2w contrast anatomical images were collected using Turbo Spin Echo (TSE) and Inversion Recovery TSE sequences, respectively, at a 0.8 x 0.8 x 1.6 mm resolution and upsampled to 0.5 mm isotropic resolution after reconstruction. All scans were acquired without sedation. Initial data preprocessing including correction for susceptibility artifacts and motion, temporal high-pass filtering, noise regression (ICA+FIX), and segmentation and generation of individual subject cortical surfaces (*70*) was provided by the dHCP group. Full details on data acquisition and initial preprocessing are reported for structural (*70*) and functional (*124*) imaging.

Adult functional MRI data were collected on a Siemens 7T Magnetom scanner with a 32-channel head coil with a single channel transmit coil at a resolution of 1.6 mm isotropic for four 16 minute runs (TR = 1 s; TE = 22.2 ms; Multiband factor = 5). Functional images were collected using a 1 s TR and multiband (factor = 5), a as resolution of 1.6 mm isotropic for four 16-minute runs. T1w structural images were collected on a custom Siemens 3T ‘Connectome’ scanner using a 32-channel head coil at a resolution of 0.7mm isotropic (TR = 2.4 s; TE = 2.14 ms). Initial data preprocessing including correction for susceptibility artifacts and motion, temporal high-pass filtering, noise regression (ICA+FIX), and segmentation and generation of individual subject cortical surfaces was provided by the HCP group (*126*, *127*). Full details on data acquisition and initial preprocessing are reported for structural and functional imaging (*126*, *127*).

### Data analysis

Analysis of Functional NeuroImages (AFNI; RRID:nif-0000-00259; Cox, 1996), SUMA (Saad and Reynolds, 2012), Freesurfer (FreeSurfer, RRID:nif-0000-00304)(*128*, *129*)(Dale et al. 1999; Fischl et al. 1999), FSL (FSL, RRID:birnlex_2067; Smith et al., 2004), Advanced Normalization Tools (ANTs) (*130*), and MATLAB (MATLAB, RRID:nlx_153890) were used for additional data processing.

### Anatomical registration to adult template and identification of probabilistic atlas

To project adult cortical visual maps into neonates, we conducted surface-based registration between each neonate’s cortical surface and the fsaverage adult surface template. Using this registration, we projected a probabilistic atlas of retinotopic maps derived from adult fMRI data (*48*) onto each neonate’s cortical surface and subsequently into their high-resolution volumetric anatomical image using AFNI’s 3dSurf2Vol function. The resulting volume was resampled to match the resolution of the functional MRI data for subsequent analysis.

The atlas registration process was validated using two complementary approaches. First, in a randomly selected subset of neonates (*n* = 30), we manually verified that the boundaries of projected visual areas aligned with anatomical landmarks typically observed in adults. Specifically, we checked that (1) the border between the dorsal and ventral components of V1 fell at the deepest point of the calcarine sulcus, (2) PHC1-2 were located in the posterior portion of the collateral sulcus, (3) IPS0 through IPS5 aligned with the superior bank of the intraparietal sulcus, and (4) V3A and V3B fell within the parieto-occipital sulcus. Second, we generated registration images for every neonate in functional space and visually confirmed that the projected visual areas aligned with cortical grey matter within regions of occipital, ventral, lateral, and dorsal cortices that correspond to expected anatomical locations in adults. In total, we analyzed 24 retinotopic areas from the retinotopic atlas: Ventral V1 (V1v), dorsal V1 (V1d), V2v, V2d, V3v, V3d, hV4, VO1, VO2, PHC1, PHC2, LO1, LO2, TO1 (MT), TO2 (MST), V3A, V3A, IPS0, IPS1, IPS2, IPS3, IPS4, IPS5, SPL1. Although TO1 and TO2 are defined based on retinotopy, they are assumed to correspond functionally to the motion-selective areas MT and MST (*48*, *55*). For consistency with broader literature, we refer to these areas as MT and MST throughout the manuscript (*99*, *131*, *132*).

To examine the large-scale organization of visual cortex, we grouped the 24 retinotopic areas based on their anatomical location: posterior occipital (V1-V3), ventral occipito-temporal (hV4, VO1-VO2, PHC1-PHC2), lateral occipito-temporal (LO1-LO2, MT, MST), and dorsal extrastriate-parietal (V3A-V3B, IPS0-IPS5) cortex. These groupings reflect established models of hierarchically organized visual pathways (*10*, *12*, *17*, *133*). Small variations in how individual areas at pathway borders (e.g., hV4) were grouped did not qualitatively change the results.

### Temporal correlation analysis

To characterize the functional organization in the neonate visual system, we computed Pearson correlation matrices between cortical regions of interest (ROIs) defined by the 24-area probabilistic retinotopic atlas in each hemisphere (48 ROIs total). The time series of each ROI was calculated by averaging the fMRI signal across all voxels within that area. For each neonate, we computed pairwise correlations between all ROIs, both within and across hemispheres. The same procedure was applied to adult participants. Individual subject matrices were aggregated by taking the median across individuals to produce group-level ROI–to-ROI correlation matrices for neonates and adults. These matrices served as the foundation for subsequent analyses, including assessments of arealization, visual pathway grouping, hierarchical organization, and the large-scale visual organization through multidimensional scaling.

### Arealization analysis

To examine whether neonates exhibited arealization, we first tested whether each area’s cross-hemisphere temporal correlations could be used to identify its homotopic counterpart (i.e. the corresponding area in the opposite hemisphere). To do so, the mean time series from each area was correlated with the time series of every voxel in the contralateral hemisphere (e.g., left V1 signal to all right hemisphere voxels). An area was considered to show arealization if the voxel wise correlations showed peak responses in an area’s homotopic counterpart (e.g., left V1 with right V1) and exhibited a stronger correlation than the anatomically adjacent (e.g., left V1 with right V2) or distal (e.g., left V1 with right MT) areas in the contralateral hemisphere.

### Multidimensional scaling of neonate visual organization

To visualize the large-scale organization of the neonate and adult visual system, we projected the ROI-to-ROI correlation matrices into two-dimensional (2D) space using multi-dimensional scaling (MDS). Prior to projection, we selected the appropriate dimensionality by applying MDS to the adult group matrix using two to five dimensions and identified the elbow point in a skree plot. Although dimensionality was determined based on the adult data, the neonate group data was also best fit using two dimensions. The resulting 2D projection showed stable layouts across random initializations for both neonates and adults. For ease of visual inspection and interpretation, the configurations were rotated so that occipital areas appeared in the bottom-left of the figure, with dorsal areas at the top and ventral areas at the bottom.

### Visual pathway grouping analysis

To assess whether visual areas were grouped according to canonical anatomical pathways, we used the 2D embeddings from each individual’s MDS analysis. For each area, we computed the Euclidean distance between its 2D position and every other area. Grouping was considered consistent with anatomical organization if a given area was, on average, closer to areas in the same pathway than to areas from different pathways. Pathway groupings were defined as follows:

- Ventral pathway: hV4, VO1, VO2, PHC1, PHC2
- Lateral pathway: LO1, LO2, MT, MST
- Dorsal pathway: V3A, V3B, IPS0–IPS5

Occipital areas (ventral and dorsal V1–V3) were treated as a separate grouping to reflect their shared position at the origin of all three pathways.

### Areal hierarchy analysis

To assess whether cortical visual areas in neonates exhibit a posterior-to-anterior hierarchical organization, we compared the spatial arrangement of areas revealed by the MDS embeddings to a canonical anatomical hierarchy defined from prior anatomical and functional studies. Each visual area was assigned a hierarchy index reflecting its known position along a given pathway:

- Ventral: V1v (1), V2v (2), V3v (3), hV4 (4), VO1 (5), VO2 (6), PHC1 (7), PHC2 (8)
- Lateral: V1d (1), V2d (2), V3d (3), LO1 (4), LO2 (5), MT (6), MST (7)
- Dorsal: V1d (1), V2d (2), V3d (3), V3A (4), V3B (5), IPS0 (6), IPS1 (7), IPS2 (8), IPS3 (9), IPS4 (10), IPS5 (11)

For each pathway, we began with the earliest area (V1v or V1d) and iteratively selected the next closest area in 2D embedding space, producing a data-driven functional ordering. We then calculated the Pearson correlation between the MDS-based ordering and the anatomically defined index values. A high correlation indicates that the functional arrangement of areas reflects the expected hierarchical structure of the visual system.

### Cortical maturity analysis

We assessed the maturity of neonatal visual cortex by comparing each area’s pattern of temporal correlations with all other areas, its “functional fingerprint,” to those of adults. Functional fingerprints refers to each area’s unique pattern of temporal correlations which can be used to identify each area (*134–136*). For each neonate, we computed the functional fingerprint of each visual area by extracting its pattern of temporal correlations with all other visual areas across both hemispheres, including the homotopic, “identical”, area in the opposite hemisphere (47 areas total). These fingerprints were derived from the previously computed ROI-to-ROI correlation matrices, where each row represents one area’s temporal correlation with every other area. We calculated fingerprints for the same visual areas in each adult participant using the same procedure. To quantify maturity, we calculated the similarity between each neonatal area’s fingerprint and the fingerprints of all adult areas. An area in the neonate was considered more mature if its functional fingerprint was most similar to the corresponding adult area, reflecting both specificity and strength of developmental correspondence.

### Eccentricity organization

We examined whether the neonate visual system exhibits evidence of an eccentricity organization. Because infants and adults were scanned at rest, and were not presented with visual stimulation, we tested eccentricity by harnessing the known foveal-to-peripheral organization across the V1 calcarine sulcus. The V1 ROI (V1v and V1d) was split into 5 ROI bands spanning the length of the calcarine sulcus (see Supplemental Figure 3). The most posterior band corresponds to foveal cortex, and therefore small eccentricities, and more anterior bands correspond to increasingly more peripheral portions of space, and therefore larger eccentricities. We then conducted cross-hemisphere seed-to-whole correlation analysis between the mean timeseries response of each ROI band with every voxel of the contralateral hemisphere. Eccentricity preferences were determined by fitting a polynomial to the 5 ROI correlations (foveal to peripheral) and taking the maximum value along the polynomial. This provides a continuous measure of a voxel’s sensitivity to eccentricity across the 5 sulcal bands.

## Acknowledgements

The authors would like to acknowledge funding from the University of Pennsylvania MindCORE fellowship and Data Driven Discovery Initiative (DDDI) fellowship awarded to V.A. as well as White hall Foundation and NIH (P50MH132642) grants awarded to M.A.

## Author Contributions

V.A. and M.A. designed the study, conducted the analysis, and wrote and edited the manuscript.

